# Harmful self-pollination drives gynodioecy in European chestnut, a self-incompatible tree

**DOI:** 10.1101/2022.08.01.502348

**Authors:** Clément Larue, Rémy J. Petit

## Abstract

- Gynodioecy is a rare sexual system in which two genders, cosexuals and females, coexist. It provides the opportunity to compare individuals having both sexual functions with individuals lacking the male function, an ideal situation to understand how sexes interact within individual plants.
- We report gynodioecy in the European chestnut, an outcrossing self-incompatible Fagaceae tree species. This finding was unexpected because gynodioecy is often interpreted as an outbreeding mechanism. To understand how female chestnuts compensate for their lack of siring capacity, we compared key female fitness components between genders and performed emasculation experiments.
- Genders have similar basal area and number of flowers but different fruit set. Following the removal of nectar-producing catkins on branches or entire trees, fruit set increased in cosexual trees but decreased in female trees.
- These results show that self-pollination impairs fruit set in cosexual trees, a likely effect of self-pollen interference caused by late-acting self-incompatibility and by early inbreeding depression. Female trees escape from self-pollen interference but continue to attract pollinators thanks to their sterile but rewarding male catkins, resulting in a much higher fruit set than cosexuals. This demonstrates that even entirely outcrossed plants can benefit from the cessation of self-pollination.

## Introduction

In angiosperms, most species have retained the ancestral hermaphrodite floral organization (Yampolsky, 1922). Starting with this sexual system well adapted for animal pollination (Sauquet *et al*., 2017), flowering plants have further evolved, resulting in an amazing variety of reproductive systems at the flower, inflorescence or whole plant level. For instance, in gynodioecy, a rare but widespread dimorphic sexual system, both cosexual (typically hermaphrodite) and unisexual (female) individuals are present. The evolutionary mechanisms leading to such gender polymorphism in plants have fascinated researchers ever since Darwin (1877). Studying these mechanisms should also help understand why the vast majority of flowering plants are cosexual (Bawa, 1984). To explore the consequences of the coexistence of both sexual functions within individuals, gynodioecious plants are better models than dioecious plants: a comparison between their two genders can help identify negative and positive interactions between sexual functions taking place within cosexual individuals (Webb, 1999). In gynodioecious species, to persist, females must compensate for their lack of male fitness (Lewis, 1941), which requires rather stringent conditions (Charlesworth & Charlesworth, 1978). This compensation, or ‘female advantage’, is the reproductive advantage of females over cosexuals due to increased life-long female fecundity through higher seed quantity, higher seed quality, or both. In recently established male-sterile mutants that lack the refined secondary characters of long-established specialist females, female advantage should be mostly attributable to the release from negative sexual interactions (Charlesworth & Charlesworth, 1978; Lloyd, 1982).

Mechanisms of female advantage can be classified in two categories, genetic and ecological (Givnish, 1982). The most cited genetic mechanism of female advantage is the outbreeding advantage, attributed to the escape from inbreeding depression in selfed offspring (Darwin, 1876; Mather, 1940; Baker, 1959; Lloyd, 1975). Other genetic mechanisms that can affect cosexuals but not females include pollen tube competition between selfed and outcrossed pollen (Shykoff, 1992) and self-pollen recognition and discrimination in the styles. The most cited ecological mechanism of female advantage is reallocation of resources to the female function from an abandoned male function (Darwin, 1877). Other ecological mechanisms of female advantage exist, such as reduction of antagonistic biotic interactions (reviewed in Ashman, 2002). In the 1980s, a heated debate took place on the nature of mechanisms driving separate sexes (Freeman *et al*., 1997), with supporters of the central importance of outbreeding advantage (Thomson & Barrett, 1981; Lloyd, 1982) being challenged by opponents emphasizing mostly ecological mechanisms (Willson, 1979; Bawa, 1980, 1982; Givnish, 1982). Forty years afterwards, we still have only a limited understanding of the relative importance of the different mechanisms (genetic or ecological) of female advantage (Dufaÿ & Billard, 2012; Pannell, 2018).

There are two main reasons why selection for outcrossing was considered by some to be the most important driver of the evolution of separate sexes. First, its principle is straightforward (Mather, 1940), and second, it has considerable predictive power. In particular, it predicts that, if hermaphrodites are partially self-fertilizing and produce lower-quality offspring because of inbreeding depression, females, but not males, will be selected. This prediction is supported by the much greater prevalence of gynodioecy over androdioecy (Lloyd, 1975; Charlesworth & Charlesworth, 1978). However, a more recent review of the origin of gender differences in gynodioecious species has shown that reduced selfing is not always the cause of female advantage (Dufaÿ & Billard, 2012). Moreover, other mechanisms than unisexuality can promote outcrossing, such as self-incompatibility (Willson, 1979). Clearly, the question of the relative importance of genetic and ecological mechanisms in the evolution of separate sexes remains open.

In females, outcrossing is enforced. However, it would be erroneous to consider that selfing avoidance is the only female advantage derived from pollen emission breakdown. Reduced seed set after self-pollination is widespread among hermaphrodite flowering plants (Burbidge & James, 1991). The cause is not merely postzygotic. In plant species with late-acting (ovarian) self-incompatibility (Seavey & Bawa, 1986), ovules are disabled by self-pollen tubes (Charlesworth, 1985; Sage *et al*., 1994; Gibbs, 2014; Johnson *et al*., 2019). Barrett *et al*. (1996) introduced the concept of ovule discounting to describe ‘*the situation where ovules are excluded from cross-fertilization because they are rendered non-functional by self-pollen tubes as a consequence of prior self-pollination*’. This represents a special case of male interference with the female function (Barrett, 2002; Duffy & Johnson, 2014). Self-pollen interference with ovules closely parallels inbreeding depression acting on embryos, seeds and seedlings: both are genetic mechanisms that ultimately limit inbreeding. In principle, if strong enough, self-pollen interference and ovule discounting could drive the evolution of separate sexes, alone or in combination with selfing and inbreeding depression. Hence, a more encompassing view would be to consider ‘self-pollination avoidance’ instead of ‘selfing avoidance’ as an explanation for any female advantage resulting from mating system change in females.

Regardless of the mechanisms involved, some circumstances can affect the success of the establishment of females in cosexual populations. First, the transmission mode of male-sterility can greatly affect female establishment. Specifically, in case of maternal inheritance of male sterility, male-sterility mutations will benefit from exclusive transmission via ovules (e.g. Lewis, 1941; Lloyd, 1975; Gouyon & Couvet, 1987). Second, the broader ecological context is also highly relevant. In particular, the establishment of females is easier when pollen is not limiting (e.g. Lloyd, 1974; Dornier & Dufaÿ, 2013; Lahiani *et al*., 2015; Stone & Olson, 2018). Third, not all sexual interactions are negative. Pollinators often prefer hermaphrodite over female flowers, due to male-biased pollinator attractiveness (Darwin, 1877; Lloyd, 1975; Wise et al., 2011; van Etten and Chang, 2014). Such positive sexual interactions can inhibit the evolution of females in cosexual populations.

Gynodioecy evolves when male-sterile mutants are able to establish. For this, the mutants must benefit not only from favourable circumstances but also from an immediate and tangible advantage (Webb, 1999). A simple but powerful method to identify mechanisms capable to bring such an instantaneous advantage is emasculation of cosexual individuals. By removing the source of pollen, we can simulate the effect of the apparition of a male-sterile mutant. Hence, relying on emasculation to investigate female advantage should help clarifying the early steps of the evolution of gynodioecy by ruling out possible pleiotropic effects of the male-sterility genes themselves as well as the effects of modifier genes selected after the establishment of sexual dimorphism (Sun & Ganders, 1986). However, one possible issue with emasculation experiments is that, by removing stamens, we remove not only the source of self-interference (pollen) but also a potential reward (pollen) as well as possible visual clues (stamens) for pollinators (e.g. Duffy and Johnson, 2011). Another possible issue with emasculation experiments that applies to hermaphrodite but not to monoecious plants is the potential effect of the experimentation on corolla development, due to corolla’s physiological interdependence with stamens (Gibson & Diggle, 1998). This illustrates the need for careful selection of model species (Charlesworth, 1993) and of emasculation controls to account for possible confounding effects. Surprisingly, given their great potential, emasculation experiments involving gynodioecious plants remain very scarce. We found only four previous studies that have explored the consequences of emasculation on female fitness in gynodioecious plants (Kikuzawa, 1989; Pettersson, 1992; Alonso & Herrera, 2001; Wang *et al*., 2021), none of which had attempted to control for an effect of the treatment on pollinator attractiveness.

Focusing on gynodioecious plants with unusual life histories or breeding systems could help understand possible limits to generalization (Freeman *et al*., 1997). Gynodioecious species tend to be herbaceous rather than woody (Caruso *et al*., 2016), hermaphrodite rather than monoecious (Lloyd, 1975), and self-compatible rather than self-incompatible (Dufaÿ and Billard, 2012). Trees are typically outcrossed (Petit & Hampe, 2006), and monoecy and self-incompatibility both promote outcrossing. Gynodioecious species with these unusual features could help explore how gynodioecy can evolve when outcrossing rates are already maximum.

In this study, to test if self-pollination avoidance is sufficient to explain the persistence of females in outbred species, we selected a long-lived Fagaceae tree, European chestnut (*Castanea sativa*). Spontaneous male-sterile variants have been reported in this species (Kaul, 1988; Soylu, 1992). Chestnut trees are monoecious, self-incompatible, and duodichogamous, all mechanisms frequently interpreted as adaptations to promote outcrossing (Lloyd & Webb, 1986; Webb & Lloyd, 1986; Bertin & Newman, 1993; Routley *et al*., 2004; Koelling & Karoly, 2007). In a previous study, we found self-fertilization rates of 5% in chestnut trees with fully developed stamens and 1% in trees with dysfunctional stamens (Larue *et al*., 2022). Even with strong inbreeding depression, this difference cannot explain the maintenance of gender dimorphism. After attempting to establish gynodioecy in the wild in European chestnut, we therefore studied if female chestnut trees have some kind of female advantage other than increased outcrossing rate. Specifically, we compared five key components of female fitness between genders in a chestnut experimental stand: basal area (a proxy for tree size), flower density, burr set, fruit set and fruit weight. Both genders harbour nectar-producing male catkins that attract pollinators; the only visible difference is that the stamens are aborted or greatly reduced in female trees (Larue *et al*., 2021a, 2022). We took advantage of this feature to assess the importance of self-pollen avoidance in an experiment where we removed male catkins not only from cosexual but also from female chestnut trees. This unique situation with female flowers associated either with fertile or with sterile but rewarding male catkins should help to disentangle the positive and negative effects of male function on female fitness.

## Materials and methods

### Chestnut floral biology

Chestnuts (*Castanea spp*.) are insect-pollinated Fagaceae tree species (Larue *et al*., 2021a; Petit and Larue, 2022). Chestnut trees are characterized by massive blooming, huge pollen production and the largest pollen:ovule ratio ever reported in plants (∼10-30 millions; Larue *et al*., 2021a). They have multiple mechanisms limiting self-pollination. First, they are monoecious, with separate male and female flowers distributed in two types of inflorescences: unisexual male catkins and bisexual catkins composed of one or two female inflorescences associated with a single male catkin (Larue *et al*., 2021a). Second, they have a complex phenology called duodichogamy, characterized by two peaks of pollen emission, which reduces the risk of self-pollination without eliminating it completely (Hasegawa *et al*., 2017). Unisexual male catkins bloom first and release huge amounts of pollen, about 97% of the total (Larue *et al*., 2021a). Non-rewarding female flowers then become receptive. Finally, male catkins from bisexual inflorescences start to emit pollen, resulting in a second much smaller pollen emission peak. Third, chestnuts are self-incompatible (Xiong *et al*., 2019).

In European chestnuts and in its hybrids, two genders can be distinguished: cosexual trees, which are fully male-fertile; and female trees, which are completely or largely male-sterile (Solignat and Chapa, 1975; Larue *et al*., 2022; Fig. **1**). These male-sterile trees have dysfunctional staminate flowers with fully aborted stamens or with short stamen filaments producing scarce amounts of mostly non-functional pollen (Bounous *et al*., 1992). Interestingly, male-sterile catkins continue to produce nectar and attract insects such as flies and beetles, but not pollen-seeking insects such as hoverflies and bees (Larue *et al*., 2021a). All flowers on a tree and all trees from a clone have the same type of male flowers (Larue *et al*., 2021b, 2022). In interspecific crosses, male sterility is expressed only in some cytoplasmic backgrounds (Bolvanský & Mendel, 1999; Sisco *et al*., 2014; Larue, 2021). In particular, in crosses between European chestnut (*C. sativa*) and Japanese chestnut (*C. crenata*), only crosses with European chestnut as mother generate male sterile individuals (Larue, 2021). However, within European chestnuts, character inheritance studies point to strict nuclear inheritance of male sterility involving a major recessive gene modified by one or more other genes (Soylu, 1992; Bolvanský & Mendel, 1999), not to nucleo-cytoplasmic inheritance, probably because cytoplasmic variation is often lacking within populations of European chestnut (Fineschi *et al*., 2000; Larue, 2021).

**Figure 1:**
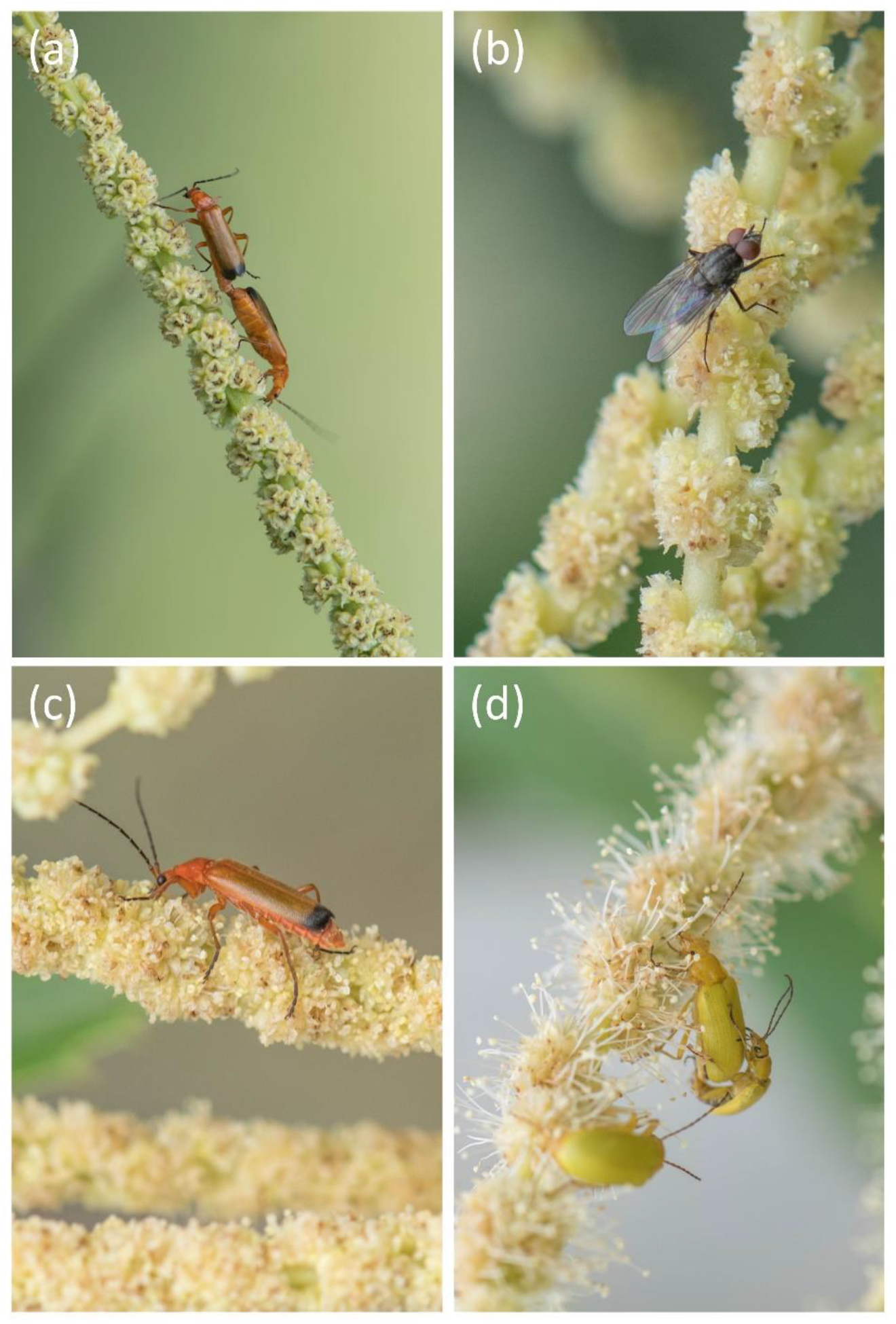
Degrees of male-sterility observed. (a) Fully male-sterile (i.e. female) tree. Most stamens are aborted and do not produce pollen. (b) Mostly male-sterile tree. Stamens do not protrude from glomerules. (c) Partly male-sterile tree. Stamens slightly protrude from glomerules. (d) Fully male-fertile (i.e. cosexual) tree. Stamen filaments are long and largely protrude from glomerules. These trees produce large amounts of pollen.

Chestnut trees, despite their long lifespan and large sizes, are good models for studying female advantage. Fruit set is easy to estimate in chestnuts. A female inflorescence is typically composed of three female flowers located side by side. The inflorescence develops into a spiny infrutescence called a burr. In each burr, each flower, if pollinated, gives a nut (a fruit with a single seed surrounded by a closed pericarp), and if not pollinated, an empty fruit. Multi-seeded nuts are very rare (< 1%, unpublished data). Hence, the proportion of developed fruits per burr should provide an indication on pollination success. Moreover, it is technically feasible to emasculate trees despite the tiny size of flowers, as male flowers are packed together into inflorescences that can be removed as one unit (Fig. **2**).

**Figure 2:**
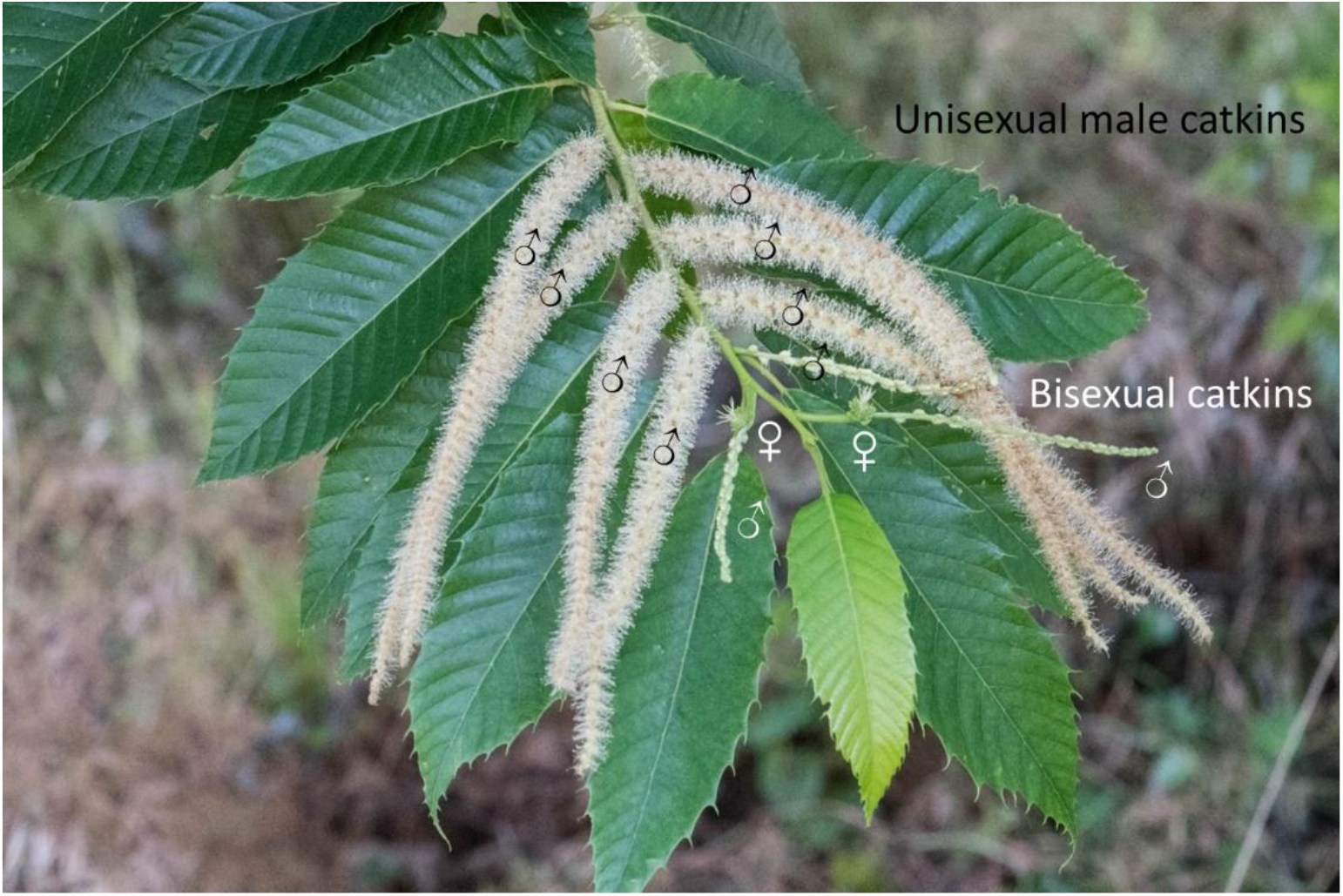
Chestnuts have two types of catkins: numerous unisexual male catkins that flower first and few bisexual catkins located at the tip of the branches that flower last.

### Gender polymorphism in chestnut forests

We visited 14 naturally regenerated populations of European chestnut, one in Spain (described in Larue, 2021) and the rest in France, and estimated the proportion of trees with different types of stamens using descriptions of Solignat & Chapa (1975) as reference (Fig. **1**). Stands were selected to be as natural as possible, avoiding ancient plantations and coppices, and focusing on regions identified as potential glacial refugia for chestnut (Krebs *et al*., 2019).

### Female advantage

#### Study site

For this work, we used the INRAE chestnut germplasm collection located in Villenave d’Ornon (44.788319 N, −0.577062 E) (Larue *et al*., 2021b). Most trees investigated come from a 3.5 ha-large experimental unreplicated orchard planted in 1990. It includes 211 grafted trees (ramets) corresponding to 83 different genotypes (clones) assigned to different chestnut species: the European chestnut (*C. sativa*; 50% of the trees), the Japanese chestnut (*C. crenata*; 9%), the Chinese chestnut (*C. mollissima*; 8%) and their interspecific hybrids, mostly *C. sativa × C. crenata* hybrids (27%). The trees are grafted on three different hybrid rootstocks: ‘Marsol’ (CA07; 56% of the trees), ‘Maraval’ (CA74; 27%), or ‘Marlhac’ (Ca118; 3%). The remaining trees (14%) grow on their own roots. Among the clones, 55 are cosexual and 28 are partly or completely male-sterile (thereafter called female), corresponding to 137 (65%) cosexual and 74 (35%) female ramets distributed rather evenly throughout the plot (Supplementary Material 1). This even-aged collection includes genotypes of forest trees as well as some clones selected for fruit production, mostly superior trees selected in nature or first generation hybrids, as selective breeding is very recent.

#### Female fitness components in the study plot and in a subset of trees

In 2019, we systematically compared several components of female fitness: basal area, fruit set and fruit weight, controlling for species status and rootstock identity. For basal area, we measured the diameter of the stem (or for multi-stemmed individuals, of each stem) at breast height and derived the cross-sectional area. To estimate fruit set, we targeted at least 30 burrs per tree. We statistically corrected fruit set to account for the possibility of premature abortion of empty burrs as explained below in the section *Statistical analyses*. To estimate fruit weight, we collected 50 developed fruits per tree and weighted them. In 2020, we studied two additional female fitness components on a subset of 18 trees, nine from each gender, belonging either to European chestnut (five clones, each with two ramets except for one clone) or to *C. sativa × C. crenata* hybrids (five clones, each with two ramets except for one clone). On 10 marked branches per tree, we counted the number of female inflorescences in the spring, the number of burrs formed in the summer, and the number of mature burrs harvested in the fall. We estimated burr set by dividing the number of burrs counted in August by the number of female inflorescences counted in June during flowering time. To investigate inter-annual fluctuations of fruit set between genders, we gathered fruit set estimates for these trees in 2018, 2019, 2020 and 2022. In view of the composition of this large collection and of the use of both female and cosexual clones in chestnut cultivation, systematic gender-biased effects of artificial selection on phenotypic traits were deemed unlikely.

### Emasculation experiment

#### Plant material

We performed the emasculation experiment in 2019 in three orchards of the INVENIO experimental station in Douville (45.019723 N, 0.614637 W). We selected six hybrid clones, three female ones (‘Bellefer’, ‘OG19’ and ‘Bouche de Bétizac’) and three cosexual ones (‘Florifer’, ‘Maraval’ and ‘Marigoule’). These trees are planted in three nearby orchards. The first orchard is composed of 8-year-old trees belonging to ‘Bellefer’, ‘OG19’, ‘Florifer’ and ‘Maraval’ clones. The second orchard is composed of 20 m-high trees belonging to ‘Bouche de Bétizac’ and ‘Marigoule’ clones. We replicated the experiment on ‘Marigoule’ clone in a third orchard, because this clone is very susceptible to cynips (*Dryocosmus kuriphilus*), which complicates fruit set measurements. Indeed, following cynips attacks, female flowers tend to proliferate, resulting in atypical inflorescences with more than three flowers that are difficult to use to estimate fruit set.

#### Modalities

To remove male flowers for the emasculation treatment, we cut all unisexual male catkins and the male part of bisexual catkins with a scissor (Fig. **2**). As these two types of catkins flower at different times, we proceeded in three steps before trees started to flower. At the first pass, in the end of May, we removed all unisexual male catkins. At the second pass, we checked if no unisexual male catkins remained and we removed the male part of bisexual catkins. At the third and last pass, we removed the male flowers of bisexual catkins that remained.

We also removed the male-sterile catkins of female clones to evaluate the decrease in insect attractiveness or possible reallocation of resources in the female function following their removal. For younger trees, for each modality, we selected five ‘OG19’ trees, six ‘Bellefer’ trees, five ‘Florifer’ trees and two ‘Maraval’ trees. Hence, there were 18 control and 18 emasculated chestnuts. We emasculated entirely each tree. For older trees, we applied the treatment to single branches. In the first orchard, we selected five ‘Bouche de Bétizac’ and five ‘Marigoule’ trees. For each tree, we selected 20 branches and used ten for each modality. We replicated these two modalities on three more ‘Marigoule’ trees from another orchard.

For young trees, we also performed partial emasculations experiments on two trees per clone for the two most abundant clones, i.e. ‘Bellefer’ and ‘Florifer’. In the ‘unisexual modality’, we removed all unisexual male catkins but left the male part of bisexual catkins. On the contrary, in the ‘bisexual modality’, we removed the male part of bisexual catkins but left all unisexual male catkins.

### Statistical analyses

#### Data analysis

We performed all analyses with R software (v3.6.6; R Core Team, 2013). The corrected fruit set was calculated with basic functions implemented in R and the violin plot is computed with ggplot2 (v3.6.3; Wickham, 2016a) and ggthemes (v4.2.4; Arnold, 2016) packages.

#### Corrected fruit set

To estimate the probability of a female flower to give a fruit, we statistically corrected fruit set to account for the possibility of premature abortion of empty burrs, i.e. burrs with no developed fruit. Not accounting for these missing empty burrs when estimating fruit set in the fall would result in an overestimation of fruit set (strong overestimation when fruit set is low, slight overestimation when it is high). Using a zero-deflated binomial distribution, we obtained unbiased estimates of fruit set for the large-scale fruit set study of 2019, using only information from burrs with at least one developed fruit (Larue *et al*., 2022). For a given tree, typical burrs developed from 3-flowers inflorescences with one, two, or three developed fruits are noted x_1_, x_2_, and x_3_ while the total number of harvested burrs is noted x_tot_. Based on the definition of multinomial distribution, a maximum likelihood estimator reflecting pollination probability 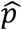 is obtained (Annex 1):

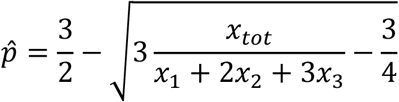

All these estimates are automatically computed for all trees in R using the *apply()* function.

In clones susceptible to cynips, inflorescences with more than three flowers can form following cynips attacks, resulting in burrs with up to nine fruits instead of the standard case with three flowers (Breisch, 1995). We excluded these rare abnormal burrs as well as even rarer cases with less than three fruits per burr from fruit set estimations.

#### Estimation of female advantage

To compare female fitness components between genders across the study plot (basal area, fruit set and fruit weight) as a function of species and rootstock, we checked ANOVA application conditions, i.e. independence, normality and homogeneity of residuals, using a Durbin-Watson test, a Shapiro test and a Bartlett test. The normality and homogeneity of the residuals were also visually inspected using the ‘Normal Q-Q’ and ‘Scale-Location’ plots. Since these conditions were not satisfied, we used non-parametric permutational analysis of variance (PERMANOVA) using *aovp()* function from lmPerm package (Wheeler & Torchiano, 2016).

We tested the differences in flower density and burr set measured in 2020 between genders and between species (European chestnut and *C. sativa* × *C. crenata* hybrids) on a subset of 18 trees using a Student *t* test (unilateral for gender effect, and bilateral for species effect).

We also tested the differences in fruit set between genders and between species during four years on the same subset of 18 trees using the same approach.

#### Emasculation experiment

We tested the significance of the difference between the modalities (control and complete emasculation) with a Fisher exact test.

## Results

### Gender polymorphism in chestnut forests

We monitored gender variation in 430 trees from 14 populations (26 to 42 trees sampled per population) (Supplementary Material 2). Male-fertile trees were more frequent in all but one population. On average, 15% of the trees were at least partly male-sterile (11% fully or mostly male-sterile and 4% partly male-sterile). Four populations were monomorphic, harbouring only male-fertile trees.

### Female advantage in chestnut

#### Basal area, fruit set and fruit weight

We found no difference in basal area between genders but significant differences among species and rootstocks (Tables 1 and 2, Supplementary Material 3). In contrast, fruit set and fruit weight measured in 2019 differed between genders but not among species or rootstocks. Fruit set was much higher in females than in cosexual trees for both European chestnut and for *C. sativa* × *C. crenata* hybrids. In contrast, fruit weight was slightly lower in female than in cosexual trees in both taxa. Fruit set and fruit weight were negatively correlated (Spearman rank-correlation coefficient: -0.15, *p*-value = 0.04).

**Table 1:** Summary table for the PERMANOVA testing of the effects of factors (gender, species and rootstock) on each female fitness component

| Fitness component | Source | Df | R Sum Sq | R Mean Sq | Iterations | Pr(Prob) |  |
| --- | --- | --- | --- | --- | --- | --- | --- |
| Basal area | Gender | 1 | 8500 | 8500 | 51 | 1.0 |  |
|  | Species | 4 | 960000 | 240000 | 5000 | <10<sup>-15</sup> | *** |
|  | Rootstock | 3 | 470000 | 160000 | 5000 | 0.006 | ** |
|  | Residuals | 202 | 8600000 | 43000 |  |  |  |
| Fruit set 2019 | Gender | 1 | 2.5 | 2.5 | 5000 | <10<sup>-15</sup> | *** |
|  | Species | 4 | 0.05 | 0.013 | 160 | 1.0 |  |
|  | Rootstock | 3 | 0.13 | 0.04 | 319 | 0.3 |  |
|  | Residuals | 167 | 5.4 | 0.03 |  |  |  |
| Fruit weight | Gender | 1 | 273 | 270 | 5000 | <10<sup>-15</sup> | *** |
|  | Species | 4 | 141 | 35 | 717 | 0.3 |  |
|  | Rootstock | 3 | 83 | 28 | 1201 | 0.4 |  |
|  | Residuals | 167 | 4000 | 24 |  |  |  |

**Table 2:** Gender effect on female fitness components

| Fitness component | #♂<sup>1</sup> | #♀<sup>1</sup> | Mean ♂<sup>2</sup> | Mean ♀<sup>2</sup> | Test<sup>3</sup> |
| --- | --- | --- | --- | --- | --- |
| Basal area (cm<sup>2</sup>) | 137 | 74 | 308 (257) | 290 (194) | ns (Permanova) |
| Flower density | 9 | 9 | 51 (34) | 67 (80) | ns (Student *t* test) |
| Burr set | 9 | 9 | 0.77 (0.15) | 0.87 (0.14) | ns (Student *t* test) |
| Fruit set | 115 | 61 | 0.57 (0.21) | 0.84 (0.11) | < 10<sup>-15</sup> (Permanova) |
| Fruit weight (g) | 115 | 61 | 10.5 (5.2) | 7.7 (4.3) | 0.0013 (Permanova) |

<sup>1</sup>Number of cosexual (♂) and female (♀) trees
<sup>2</sup>Mean value for cosexual and female trees (standard deviation)
<sup>3</sup>Difference significance (test performed)

#### Flower density and burr set

Flower density was highly variable among trees and there was no significant difference between genders or between species (Tables 2 and 3, Supplementary material 4). Similarly, there were no difference in burr set between genders nor between species in our sample.

**Table 3:** Fruit set measured during four years and details for 2020

| Gender | Fruit set |  |  |  | Details for 2020 |  |  |
| --- | --- | --- | --- | --- | --- | --- | --- |
|  | 2018 | 2019 | 2020 | 2022 | #♀-inflo<sup>1</sup> | #Burrs<sup>2</sup> | Burr set<sup>3</sup> |
| Female | 0.83 | 0.77 | 0.69 | 0.86 | 67.1 | 59.9 | 0.86 |
| Cosexual | 0.64 | 0.49 | 0.34 | 0.34 | 50.8 | 35.9 | 0.77 |
| **Ratio F/C** | **1.30** | **1.56** | **2.01** | **2.54** | **1.32** | **1.67** | **1.12** |
| *t-test* | 0.003 | 0.002 | 0.004 | <10<sup>-6</sup> | ns | ns | ns |

<sup>1</sup>Mean number of female inflorescences per tree (10 branches per tree)
<sup>2</sup>Number of burrs counted in August
<sup>3</sup>Burr set calculated by dividing the number of burrs by the number of female inflorescences

#### Fruit set across years

For female trees, average fruit set fluctuated during the four studied years, ranging from 0.69 in 2020 to 0.83 in 2018 (Table 3, Fig. **3**, Supplementary material 4). For cosexual trees, average fruit set fluctuated more drastically, ranging from 0.34 in 2022 to 0.64 in 2018. During all four fruiting episodes, fruit set was higher in females: the female/cosexual fruit set ratio ranged from 1.3 in 2018 and 1.6 in 2019, to 2.0 in 2020 and 2.5 in 2022.

**Figure 3:**
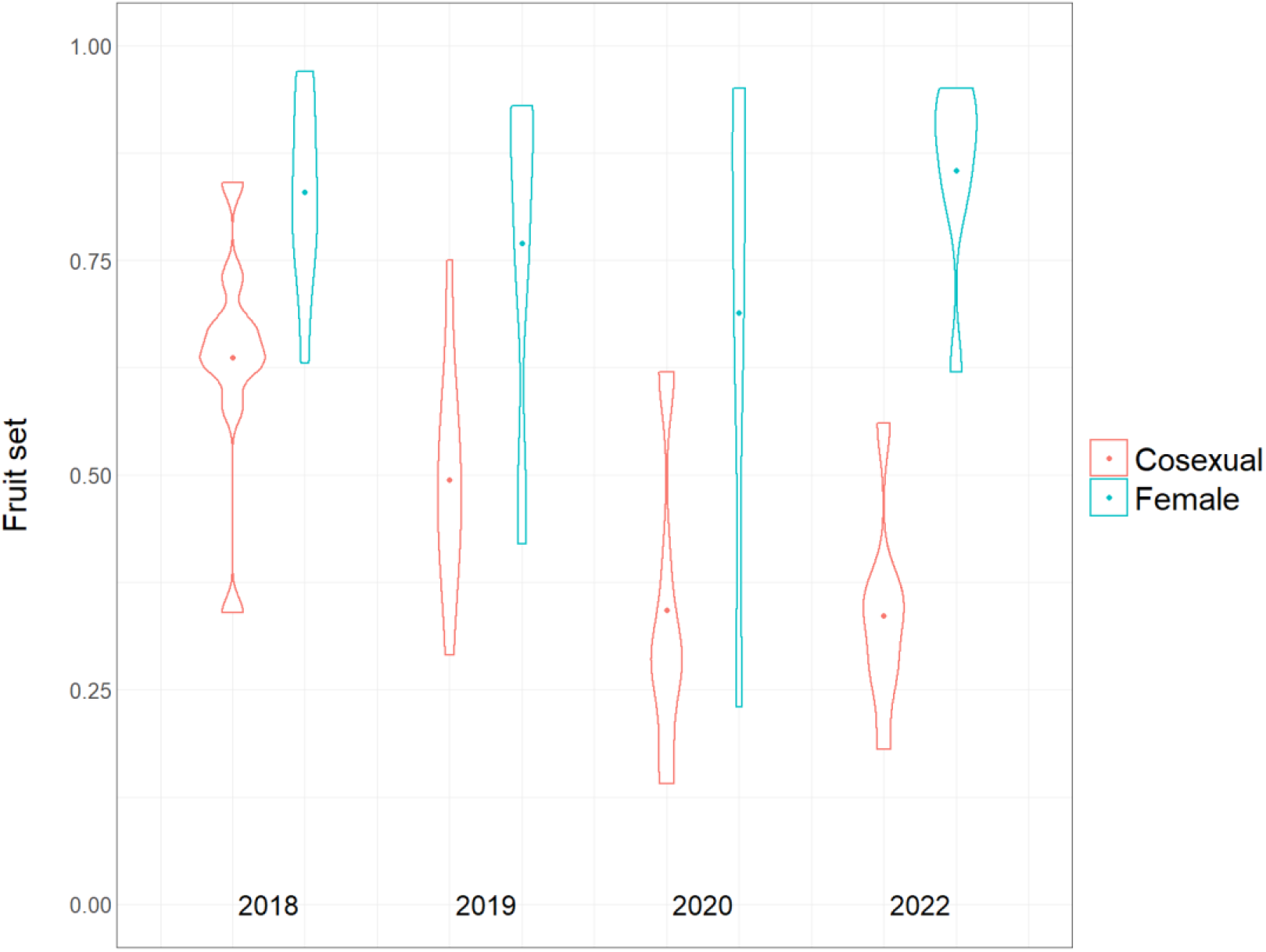
Inter-annual variation in fruit set in cosexual and female chestnut trees.

### Emasculation experiment

For older trees, the emasculation treatment resulted in a significant increase in fruit set compared to controls for the cosexual clone (Table 4). Fruit set in control trees was 52% whereas it was 67% for emasculated trees (Fisher exact test, *p* < 10^−5^). In contrast, the emasculation treatment resulted in a significant decrease in fruit set compared to control for the female clone. Fruit set in control trees was 95% whereas it was 85% for emasculated trees (*p* < 10^−7^).

**Table 4:** Fruit set of control and emasculated adult trees

| Gender<sup>1</sup> | Clone | Modality | #replicates | Mean (sd) | Fisher<br>exact test |
| --- | --- | --- | --- | --- | --- |
| F | Betizac | Control | 5 ramets × 10 branches | 0.95 (0.04) | $10^{-6}$ |
| F | Betizac | Emasculated | 5 ramets × 10 branches | 0.85 (0.07) | |
| C | Marigoule | Control | 8 ramets × 10 branches | 0.52 (0.08) | $10^{-5}$ |
| C | Marigoule | Emasculated | 8 ramets × 10 branches | 0.66 (0.07) | |

<sup>1</sup>F: Female (male-sterile); C: cosexual (male-fertile, monoecious)

Table 4: Fruit set of control and emasculated adult trees
| Gender<sup>1</sup> | Clone | Modality | #replicates | Mean (sd) | Fisher exact test |
| --- | --- | --- | --- | --- | --- |
| F | Betizac | Control | 5 ramets × 10 branches | 0.95 (0.04) | 10<sup>-6</sup> |
| F | Betizac | Emasculated | 5 ramets × 10 branches | 0.85 (0.07) |  |
| C | Marigoule | Control | 8 ramets × 10 branches | 0.52 (0.08) | 10<sup>-5</sup> |
| C | Marigoule | Emasculated | 8 ramets × 10 branches | 0.66 (0.07) |  |

<sup>1</sup>F: Female (male-sterile); C: cosexual (male-fertile, monoecious)

For younger trees, which were entirely emasculated, the emasculation treatment resulted in an increase in fruit set compared to controls for both cosexual clones (Table 5). For ‘Florifer’, fruit set in control trees was 46% whereas it was 63% for emasculated trees, a significant increase (*p* < 10^−10^). For ‘Maraval’, fruit set of control trees was 73% compared to 78% for emasculated trees, a non-significant increase (*p* > 0.16). In contrast, the emasculation treatment resulted in a significant decrease in fruit set compared to control for one of the two studied female clones. For ‘Bellefer’, fruit set in control trees was 91% whereas it was 81% for emasculated trees, a significant decrease (*p* < 10^−5^). However, for ‘OG19’, fruit set in control trees was 77% whereas it was 82% for emasculated trees, a non-significant increase (*p* > 0.8).

**Table 5:** Fruit set of control and emasculated young trees

| Gender<sup>1</sup> | Clone | Modality | #trees | Mean (sd) | Fisher exact test |
| --- | --- | --- | --- | --- | --- |
| F | Bellefer | Control | 5 | 0.91 (0.04) | 10<sup>-5</sup> |
| F | Bellefer | Emasculated | 5 | 0.81 (0.11) |  |
| F | OG19 | Control | 5 | 0.77 (0.14) | ns |
| F | OG19 | Emasculated | 5 | 0.82 (0.05) |  |
| C | Maraval | Control | 2 | 0.73 (0.09) | ns |
| C | Maraval | Emasculated | 2 | 0.78 (0.06) |  |
| C | Florifer | Control | 6 | 0.46 (0.14) | 10<sup>-10</sup> |
| C | Florifer | Emasculated | 6 | 0.63 (0.14) |  |

<sup>1</sup>F: Female (male-sterile); C: cosexual (male-fertile, monoecious)

We performed partial emasculations experiments on two clones (Supplementary Table 5). For ‘Florifer’, a cosexual clone, fruit set increased compared to the control after removing all male catkins or only catkins from unisexual inflorescences. On the other hand, when we removed catkins from bisexual inflorescences, fruit set decreased compared to the control. For ‘Bellefer’, a female clone, fruit set following partial emasculations were intermediate between the control (higher fruit set) and the complete emasculation treatment (lower fruit set), with more reduced fruit set after removing male catkins from bisexual inflorescences.

## Discussion

Finding gynodioecy in European chestnut was unexpected. While dioecy and monoecy are particularly frequent in trees, stable gynodioecy is rare (Olson *et al*., 2016; but see Steyn & Robbertse, 1990; Ellis & Sedgley, 1993; Gibson & Diggle, 1998; Penagos Zuluaga *et al*., 2020). Moreover, gynodioecy is unknown in Fagaceae, a widespread species-rich family (Petit *et al*., 2013). Chestnut is an ecologically and economically important genus cultivated for its nutritious nuts. The fact that gynodioecy had remained unnoticed in chestnuts seems therefore surprising. In gynodioecious plants, the average female frequency is 25% (Varga & Soulsbury, 2020). If gender variation is studied in few populations, evidence for gynodioecy might be missed (Dufaÿ *et al*., 2014; Caruso *et al*., 2016). Our survey illustrates this point, as we found a rather low (15%) but variable (0-53%) proportion of females, so that half of the populations have less than 10% of female trees. We also found some partly male-sterile trees, a frequent feature in gynodioecious species (Koelewijn & van Damme, 1996; Schultz, 2002). Because spontaneous male sterile mutants have been reported in other chestnut species (Kaul, 1988), exploring gender variation in other chestnut species would be worthwhile.

To explore female advantage in chestnut, we investigated for the first time in a tree several key female fitness components covering the complete life cycle (Philipp, 1980). Using a grafted clonal collection established for conservation purposes, we found no difference in basal area and flower production between genders. In grafted trees, the rootstock partly controls a number of phenotypic traits, including growth, but scions are known to influence tree canopy height in chestnut (Camisón *et al*., 2023), so the lack of gender effect on scion growth is remarkable. This finding suggests that there is no significant reallocation of resources to the female function from the lost male function at this stage. We also found that females have lighter, not heavier, fruits than cosexual trees. Fruit weight is negatively correlated with fruit set, a likely effect of competition for resources between fruits within female inflorescences (Larue, 2021). Therefore, females do not appear to reallocate significant resources to fruit growth. Long-lived perennial plants need to allocate more resources than short-lived plants for the maintenance of vegetative structures, so intersexual competition for resources plays a relatively smaller role in long-lived than in short-lived plants (Obeso, 2002). Furthermore, reallocation of resources from male toward female function is necessarily limited in gynodioecious trees because of the higher cost for female than for male reproduction in woody plants (Obeso, 2002; Thomas, 2011).

The single female advantage identified in chestnut is increased fruit set. This advantage is large and holds for both European chestnut and for its hybrids. Japanese and Chinese chestnuts, for which evidence for gynodioecy is lacking, had fruit sets similar to those of cosexual European chestnut trees and hybrids. To our knowledge, gender effect on fruit set has not been investigated before. However, chestnut cultivation in Europe relies heavily on female clones (Furones-Pérez & Fernández-López, 2009) and some researchers have noticed that female chestnuts tend to have higher yields (e.g. Pereira-Lorenzo & Ramos-Cabrer, 2004). We observed higher and more stable fruit set in females than in cosexual trees during four years. We tentatively attribute the less stable fruit set of cosexual trees to Jensen’s inequality (Jensen, 1906; Garibaldi *et al*., 2011): as fruit set decelerates with increased pollen receipt, the deficit of a bad pollination episode is necessarily higher than the benefit of a good pollination episode. This asymmetrical response is more pronounced at lower pollination rates where small changes translate into large differences. What matters for long-term female maintenance is the difference in geometric lifetime fitness between genders (Eckhart, 1992). Hence, episodes in which cosexual individuals have particularly reduced fruit set compared to females will accentuate long-term female advantage. Altogether, our experimental data support the hypothesis that female advantage is sufficient for female maintenance, as it is close to the two-fold threshold needed in the case of nuclear inheritance of male sterility (Lewis, 1941).

The emasculation experiments comfort the hypothesis that increased fruit set is the main female advantage in chestnut. When we removed male catkins (the sources of self-pollen) in cosexual trees, fruit set increased. In contrast, when we removed male-sterile catkins from female trees, fruit set decreased or remained unchanged. These contrasting effects suggest that the presence of pollen in anthers is decisive. On the contrary, reallocation of resources from the male catkins following emasculation is unlikely, as it should affect both female and cosexual trees. We thus attribute the opposite effects of emasculation on fruit set in the two genders to decreased self-pollination and associated ovule and seed discounting in emasculated cosexual trees and to reduced attractiveness to pollinators in ‘emasculated’ female trees. Together, these findings suggest that, in cosexual trees, the positive effect of emasculation caused by the elimination of the source of self-pollination overcomes the negative effect of decreased pollinator attractiveness caused by the elimination of the rewarding catkins. In a previous emasculation experiment involving Chinese chestnut, Zhao and Liu (2009) described increased fruit yield by up to 39%, indicating that self-pollination also negatively affects fruit production in this species not known to be gynodioecious. The finding that a single main fitness component (fruit set) determines female advantage and that this advantage can be reproduced in cosexual trees by removing the source of pollen is compatible with the hypothesis of a relatively recent origin of gynodioecy in European chestnut. Other studies have reported increased seed set following emasculation of self-incompatible hermaphrodite plant species (e.g., Waser & Price, 1991; Vaughton & Ramsey, 2010; Duffy *et al*., 2013, 2021), suggesting that self-pollen often interferes with female fitness in outcrossing plants.

What is the cause of the large difference in fruit set between genders? Self-pollination in cosexual chestnut trees is massive, a likely consequence of huge pollen production and frequent geitonogamy in this large mass-flowering tree. In the study site, we previously estimated, using a spatially explicit mixed mating model, that the self-pollination rate is 74% (Larue et al. 2022). Hasegawa *et al*. (2009) found even higher values of self-pollination in Japanese chestnut by genotyping pollen grains directly collected on stigmas. The model-based estimate of the proportion of ovules that are eventually self-fertilised is 48%, as cross-pollen is somehow favoured over self-pollen. However, according to the model, 95% of the remaining self-fertilized ovules abort, meaning that 46% of ovules are lost in cosexual trees, a very large proportion (Larue et al. 2022). Cytological studies and pollination experiments have revealed both prezygotic late-acting self-incompatibility and early-acting inbreeding depression in chestnut (Xiong et al. 2019). Self-pollen tubes grow well in the styles but many ovules abort when pollen tubes reach the ovary. If self-fertilization nevertheless takes place, the resulting embryos stop growing before a mature seed is formed due to early inbreeding depression (Xiong et al. 2019). Early inbreeding depression is an important but often neglected component of inbreeding depression (Lande *et al*., 1994), particularly in outcrossing species (Husband & Schemske, 1996). An obvious advantage of early elimination of self-pollinated ovules and selfed embryos is the prevention of wasteful provisioning of low-quality progeny (Johnson *et al*., 2019; Larue *et al*., 2022). Hence, in chestnuts, gynodioecy should not be considered as a superfluous, redundant outcrossing mechanism (Baker, 1963; Krohne *et al*., 1980). Little attention has been given so far to self-pollen interference as a source of female advantage (but see Kikuzawa, 1989). We suggest that it is time to reconsider the evidence, as late-acting self-incompatibility is widespread, especially in woody plants (Seavey & Bawa, 1986; Sage *et al*., 1994; Gibbs, 2014; Johnson *et al*., 2019).

We used female trees as ‘natural controls’ in the emasculation study, removing their rewarding but sterile male catkins. This decreased their fruit set, showing that even sterile male catkins promote pollination. Our pilot partial emasculation experiment further suggests that it is the late-flowering male catkins physically associated with female inflorescences that attract insects close to female flowers (van de Pijl, 1978; Larue et al., 2021a). We previously showed that insect visits to female trees are reduced compared to visits to cosexual trees, but only for insects feeding on pollen, such as syrphid flies and bees (Larue *et al*., 2021a). However, the scarcity of these insects on female trees has no adverse consequence for pollination because they fail to visit female flowers. In contrast, beetles and calyptrate flies, the putative pollinators of chestnut, visit both genders equally. Hence, in chestnuts, the breeding system (monoecy) and main reward type (nectar) is not an obstacle to the evolution of gynodioecy.

The emasculation experiments involved either single branches or entire trees, resulting in both cases in increased fruit set in cosexual trees. Beetles are particularly abundant in chestnut groves, walking back and forth on the crown, presumably bringing much self-pollen from nearby male catkins on female inflorescences (Larue *et al*., 2021a). By emasculating branches rather than entire trees, we reduce self-pollination brought about by the behaviour of these insects, not by the behaviour of insects flying among branches of the same tree, such as flies. This suggests that insects that tend to walk on inflorescences rather than to fly from one place to another are major causes of self-pollination. Greater abundance of those pollinators responsible for insect-assisted geitonogamous mating could widen the fitness gap between genders and further promote gynodioecy.

In European chestnut, female and cosexual trees are equally outcrossed, ruling out outbreeding advantage as possible cause of female advantage. According to Baker (1959), ‘*one might expect that in any hermaphrodite taxon where an incompatibility system is already established (or even a strongly effective outbreeding system of some other sort) there will be little likelihood of dioecism arising through direct natural selection. Any exceptions to this rule would be in the nature of unstoppable accidents with special causes*’. Our study, focusing precisely on one of these exceptions, illustrates that the ‘special causes’ mentioned by Baker are no accidents after all. Rather, they involve important early acting genetic mechanisms of plant reproduction. Because females avoid self-pollination and its deleterious consequences (Charlesworth, 1993), they are more effectively outcrossed than cosexual individuals, if outcrossing success is measured in numbers of outcrossed progenies rather than in proportion of outcrossed progenies. Considering ‘self-pollination avoidance’ rather than ‘selfing avoidance’ as key female advantage could clarify the debate on the relative importance of genetic versus ecological mechanisms for the evolution of unisexuality (Freeman *et al*., 1997). The world of possibilities for the evolution of female advantages is vast but some pathways might be more important than others, compounded by plant species life histories.

## Acknowledgements

We thank Teresa Barreneche (chestnut germplasm collection administrator) for her invaluable support, and Catherine Bodénès for her collaboration and for allowing us to report the first results of her study on gender variation in natural chestnut populations from southern France. This study was possible thanks to the continuous management of the experimental plots by INRAE experimental unit Vigne Bordeaux (UEVB) and by Xavier Capdevielle of UMR Biogeco. We thank Yannick Mellerin, Céline Lalanne, Chantal Helou Bagate and our many great students for their invaluable help in the field. We also thank the INVENIO chestnut team and the numerous seasonal workers who helped with the emasculation experiment. Many thanks to Olivier Lepais and to Etienne K. Klein for developing a fruit set model and to Josefina Fernández López for sharing her knowledge on chestnut reproduction and guiding us in the field. Mathilde Dufaÿ comments helped improve an earlier version of this manuscript.

This paper was part of the PhD and postdoc of CL. The Cifre thesis (Conventions Industrielles de Formation par la Recherche) was supported by the ANRT (Association Nationale de la Recherche de la Technologie, convention Cifre N° 2018/0179). It was funded by Invenio, the Région Nouvelle-Aquitaine (Regina project N° 22001415-00004759 and chestnut pollination project N° 2001216-00002632), by INRAE (Institut National de Recherche pour l’Agriculture et l’Environnement) and by ANR (project ANR-21-PRRD-0008-01).

## Author contributions

CL, RJP and Biogeco colleagues estimated fruit set in the INRAE chestnut germplasm collection. CL and Invenio colleagues carried out the emasculation experiments in Invenio orchards. CL and RJP estimated fruit set in INVENIO orchards. CL corrected fruit set thanks to the statistical correction developed by Etienne K. Klein. CL and RJP carried out statistical analyses. CL wrote the first draft of this article and RJP revised it.

## Data availability

The data that support the findings of this study are openly available at: https://entrepot.recherche.data.gouv.fr/dataset.xhtml?persistentId=doi:10.57745/MOTPXM

They can be cited as: Larue, Clément, 2022, “DATA: Self-interference and female advantage in chestnut”, https://doi.org/10.57745/MOTPXM, Recherche Data Gouv, V1.

The data on gender variation that support the findings of this study are openly available at: https://entrepot.recherche.data.gouv.fr/dataset.xhtml?persistentId=doi:10.57745/LFZFT2

They can be cited as: Bodénès, Catherine, 2023, “Conservation des ressources génétiques de châtaignier”, Recherche Data Gouv, V1.

## Competing interest

The authors declare that they have no known competing financial interests or personal relationships that could have appeared to influence the work reported in this paper.

## Supplementary Materials

**Supplementary material 1:**
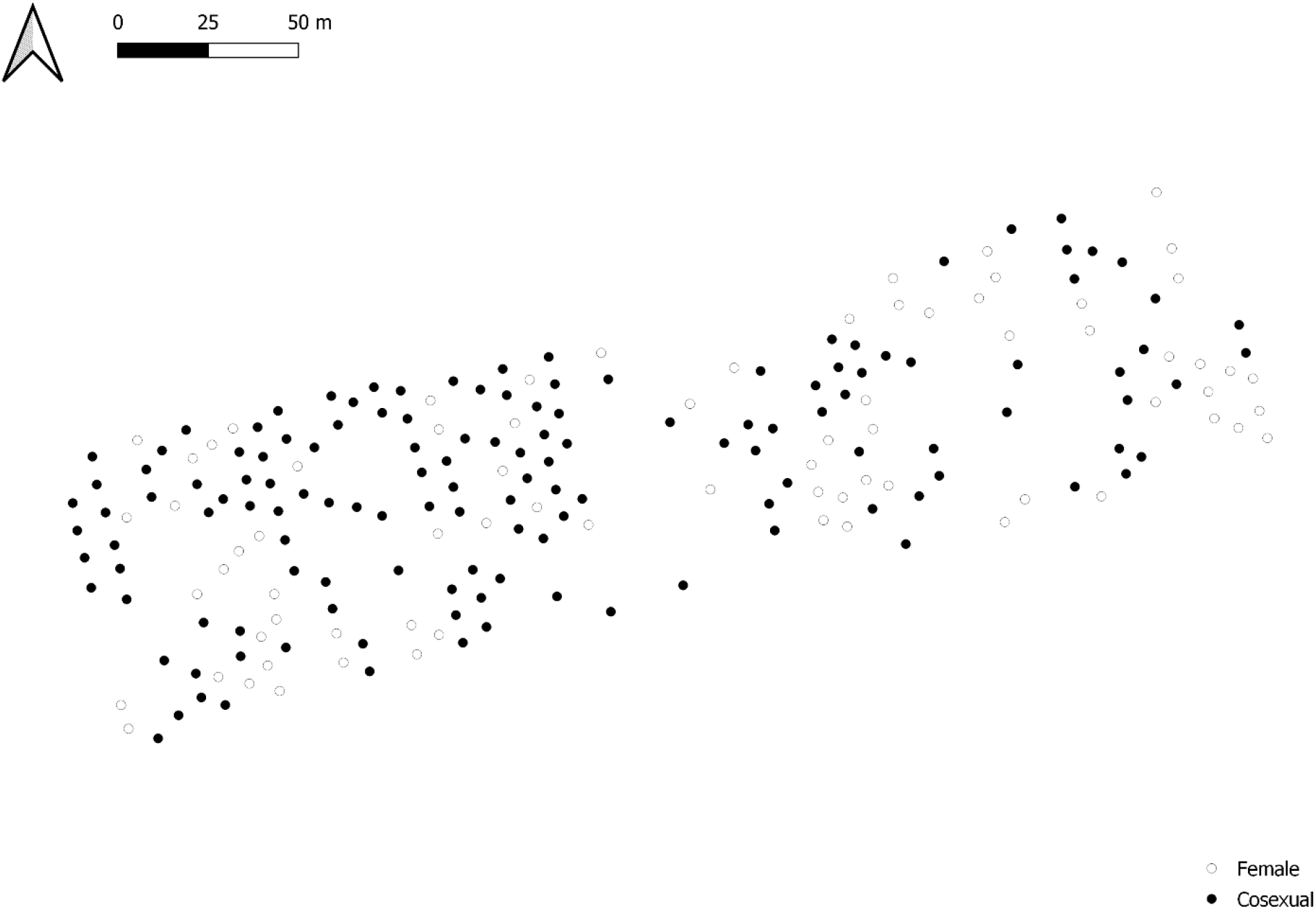
Map of the chestnut experimental plot 1.

**Supplementary material 2:**
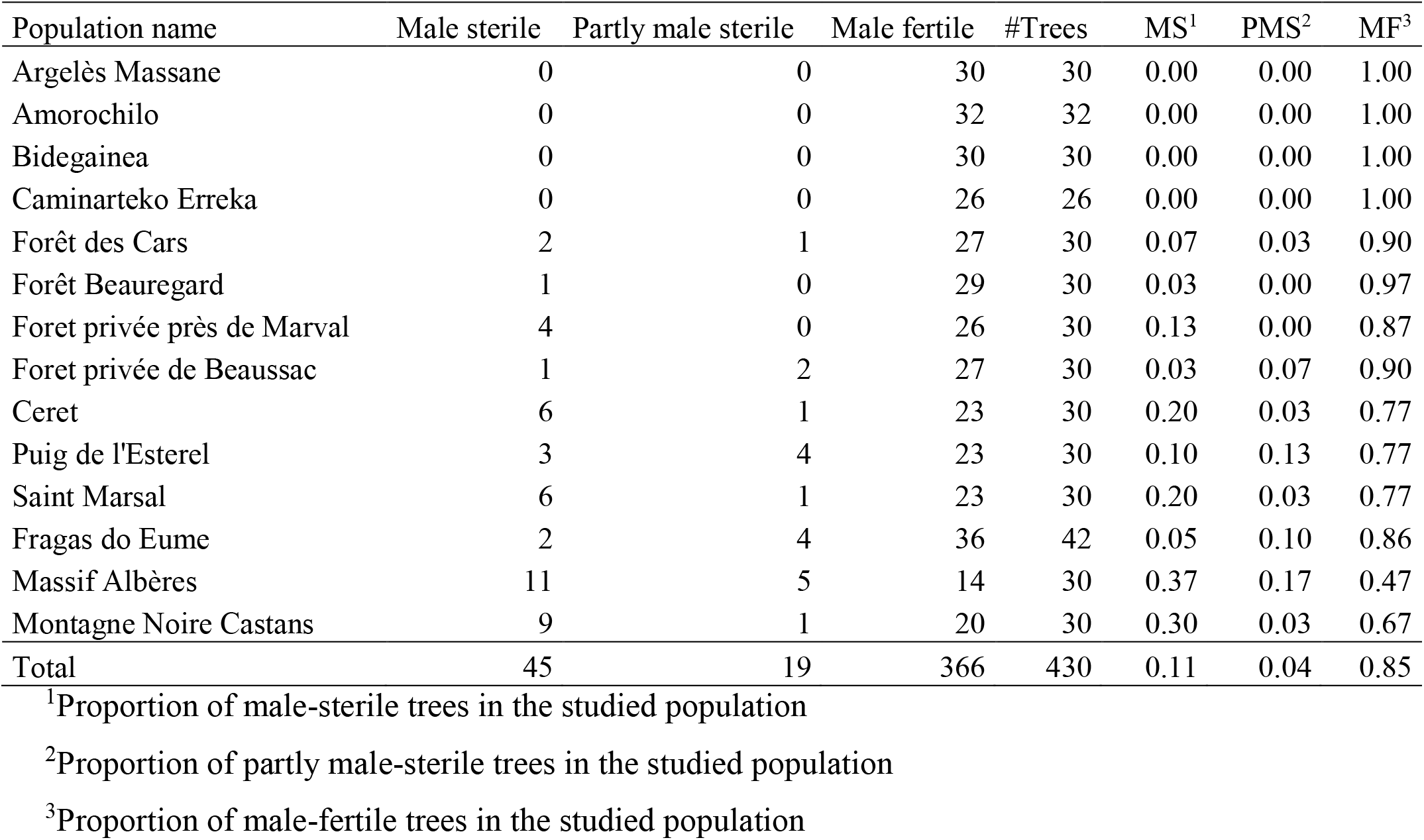
Estimation of gender polymorphism in chestnut forests

**Supplementary material 3:**
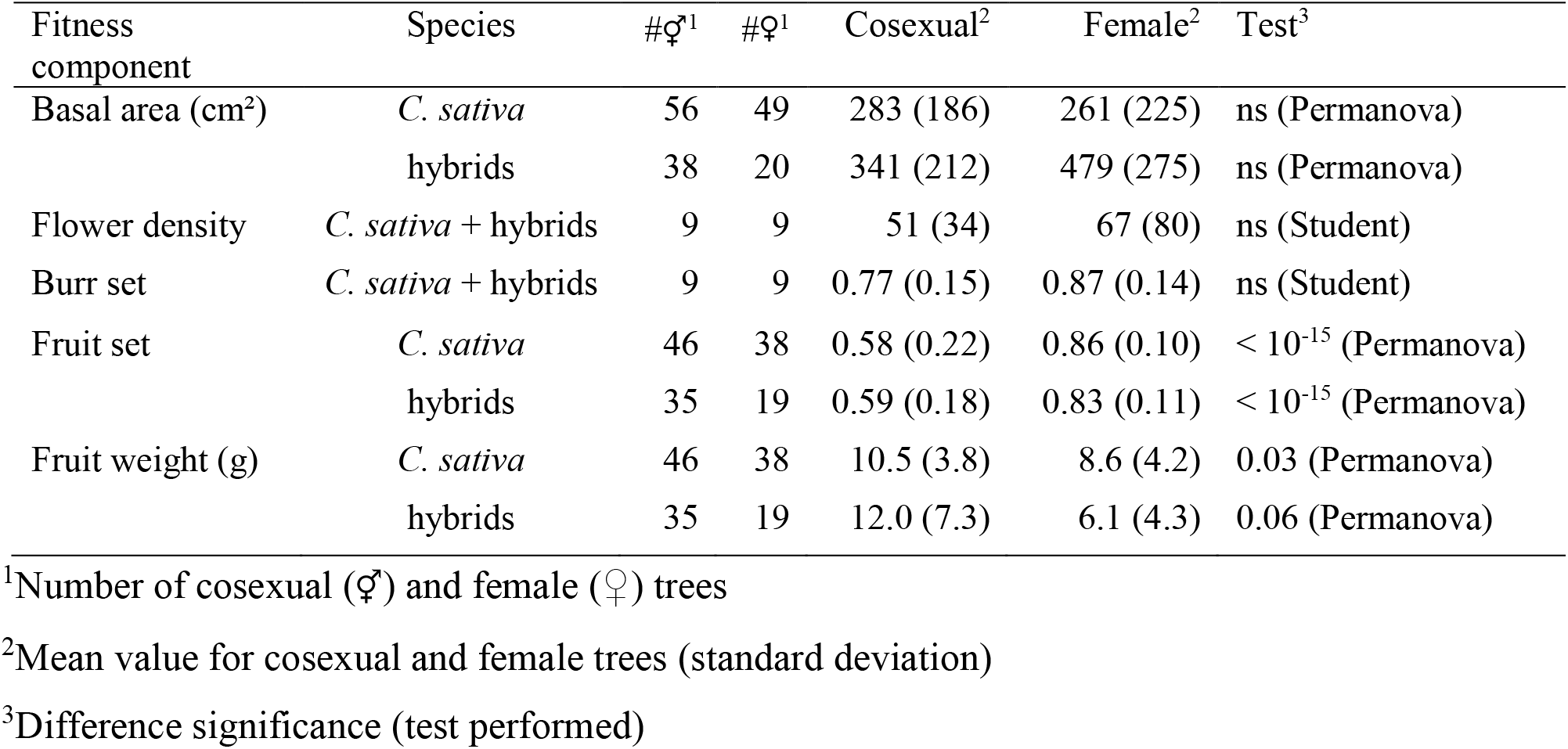
Gender effect on female fitness components by species

**Supplementary material 4:** Fruit set measured during four consecutive years and details for 2020

|  |  |  |  | Fruits set |  |  |  | Details for 2020 |  |  |
| --- | --- | --- | --- | --- | --- | --- | --- | --- | --- | --- |
| Clone | Ramet<sup>1</sup> | Species<sup>2</sup> | Gender<sup>3</sup> | 2018 | 2019 | 2020 | 2022 | #♀-inflo<sup>4</sup> | #Burrs<sup>5</sup> | Burr set<sup>6</sup> |
| Dauphine | A_64_C | S | F | 0.63 | 0.42 | 0.23 | 0.62 | 42 | 33 | 0.79 |
| Dauphine | E_76_I | S | F | 0.76 | 0.93 | 0.67 | 0.85 | 5 | 3 | 0.60 |
| Précoce des Vans | E_40_R | S | F | 0.96 | 0.92 | 0.95 | 0.95 | 57 | 46 | 0.81 |
| Précoce des Vans | E_72_S | S | F | 0.89 | 0.82 | 0.92 | 0.82 | 25 | 22 | 0.88 |
| Merle | A_45_E | S | F | 0.85 | 0.50 | 0.26 | NA |  |  |  |
| Bétizac | A_58_O | SC | F | 0.97 | 0.92 | 0.89 | 0.94 | 16 | 17 | 1.06 |
| Bétizac | E_80_E | SC | F | NA | 0.92 | 0.94 | NA | 42 | 40 | 0.95 |
| Marlhac | A_66_Q | SC | F | 0.80 | 0.75 | 0.62 | 0.91 | 254 | 230 | 0.91 |
| Marlhac | E_73_D | SC | F | 0.77 | 0.75 | 0.72 | 0.89 | 96 | 88 | 0.92 |
| 381 | E_43_O | S | C | 0.63 | 0.40 | 0.28 | 0.18 | 46 | 40 | 0.87 |
| 381 | E_52_J | S | C | 0.67 | 0.47 | 0.29 | 0.28 | 32 | 28 | 0.88 |
| Despont n3 | A_65_F | S | C | 0.58 | 0.47 | 0.39 | 0.35 | 49 | 35 | 0.71 |
| Despont n3 | E_78_E | S | C | 0.67 | 0.57 | 0.62 | 0.35 | 15 | 13 | 0.87 |
| Maridonne | A_60_Q | SC | C | 0.64 | 0.51 | 0.28 | 0.56 | 128 | 63 | 0.49 |
| Maridonne | E_50_P | SC | C | 0.73 | 0.75 | 0.62 | NA | 63 | 47 | 0.75 |
| Marigoule | A_42_N | SC | C | 0.34 | 0.29 | 0.17 | 0.27 | 44 | 42 | 0.95 |
| Marigoule | A_44_Q | SC | C | 0.63 | 0.59 | 0.29 | 0.36 | 29 | 19 | 0.66 |
| Vignols | A_55_S1 | SC | C | 0.84 | 0.40 | 0.14 | NA |  |  |  |
|  |  |  | Female | 0.83 | 0.77 | 0.69 | 0.86 | 67.1 | 59.9 | 0.86 |
|  |  |  | Cosexual | 0.64 | 0.49 | 0.34 | 0.34 | 50.8 | 35.9 | 0.77 |
|  |  |  | **Ratio F/C** | **1.30** | **1.56** | **2.01** | **2.54** | **1.32** | **1.67** | **1.12** |
|  |  |  | *t-test* **Gender** | **0.003** | **0.002** | **0.004** | **<10<sup>-6</sup>** | **ns** | **ns** | **ns** |
|  |  |  | **Species** | **ns** | **ns** | **ns** | **ns** | **ns** | **ns** | **ns** |

<sup>1</sup>Identification code of each studied tree (ramet)
<sup>2</sup>Species: *C. sativa* (S) / *C. sativa* × *C. crenata* (SC)
<sup>3</sup>Gender: female (F) / cosexual (C)
<sup>4</sup>Number of female inflorescences per tree (10 branches per tree)
<sup>5</sup>Number of burrs per tree counted in August (10 branches per tree)
<sup>6</sup>Burr set calculated by dividing the number of burrs by the number of female inflorescences

**Supplementary material 5:** Fruit set following partial and complete emasculation

| Gender<sup>1</sup> | Clone | Modality | #trees<sup>2</sup> | Mean |
| --- | --- | --- | --- | --- |
| F | Bellefer | Control | 5 | 0.91 |
| F | Bellefer | Bisexual | 2 | 0.83 |
| F | Bellefer | Unisexual | 2 | 0.87 |
| F | Bellefer | Complete | 5 | 0.81 |
| C | Florifer | Control | 6 | 0.46 |
| C | Florifer | Bisexual | 2 | 0.39 |
| C | Florifer | Unisexual | 2 | 0.49 |
| C | Florifer | Complete | 6 | 0.62 |

<sup>1</sup>Gender of studied trees: F = Female, C = cosexual
<sup>2</sup>Number of studied trees in each modality

## Notes

https://doi.org/10.57745/MOTPXM

https://entrepot.recherche.data.gouv.fr/dataset.xhtml?persistentId=doi:10.57745/LFZFT2

### Competing Interest Statement

The authors have declared no competing interest.

### Summary of Updates

This version of the manuscript has been entirely revised: in particular, we extended the scope of the paper, added new results and rewrote the Introduction and Discussion.

